# Prenatal aromatase inhibition alters postnatal immune function in domestic chickens (*Gallus gallus*)

**DOI:** 10.1101/249748

**Authors:** J.W. Simkins, F. Bonier, Z.M. Benowitz-Fredericks

**Affiliations:** Bucknell University, 701 Moore Avenue, Lewisburg, Pennsylvania, USA; Queen’s University, 99 University Ave, Kingston, ON K7L 3N6, Canada

**Author notes:** Current address: Montana State University, Bozeman, Montana 59717 (J.W. Simkins). Corresponding author: Email addresses: JWS FB ZMBF.

**Keywords:** Testosterone, Estradiol, Fadrozole, Immunity, Development, Birds

## Abstract

In birds, exposure to testosterone during embryonic development can suppress immune function; however, it is unclear whether this is caused by direct stimulation of androgen receptors. Estradiol is synthesized from testosterone by the enzyme aromatase, and this conversion is a necessary step in many signaling pathways that are ostensibly testosterone-dependent. Many lines of evidence in mammals indicate that estradiol can affect immune function. We tested the hypothesis that immunosuppressive effects of avian *in ovo* testosterone exposure are mediated by conversion to estradiol by aromatase, using Fadrozole to inhibit aromatization of endogenous testosterone during a crucial period of embryonic immune system development in domestic chickens (*Gallus gallus*). We then measured total IgY antibody count, response to PHA challenge, mass of thymus and bursa of Fabricius, and plasma testosterone post-hatch on days 3 and 18. We predicted that if immunomodulation by testosterone is dependent on aromatization, then Fadrozole treatment would lead to elevated immune activity by inhibiting estrogen production. Conversely, if testosterone inhibits immune function directly by binding to androgen receptors, then Fadrozole treatment would likely not alter immune function. Fadrozole treated birds had decreased day 3 plasma IgY antibody titers but there was a strong trend towards increased day 18 thymic mass. Furthermore, Fadrozole treatment generated a positive relationship between testosterone and thymic mass in males, and tended to increase day 18 IgY levels for a given bursal mass in females. There was no effect on PHA response, bursal mass, or plasma testosterone at either age. Overall, Fadrozole treated birds tended to have elevated indicators of immune function, implicating aromatization as a relevant pathway through which developmental exposure to testosterone can affect immunity.

## 1. INTRODUCTION

The vertebrate immune system is responsible for defending the body from a variety of external threats and must be tightly regulated to optimize protection from disease. Investment in immune defenses begins as early as oocyte formation, when the mother deposits maternal antibodies, which will form the basis of embryonic immunity, into the egg (Ulmer-Franco, 2012). Of the multiple environmental and genetic factors that influence immune function, differential hormonal exposure is a prominent source of immunological variability (Olsen and Kovacs, 1996). While variation in hormone levels can transiently affect immune function in adults, hormone-mediated actions during development have the potential to permanently alter the trajectory of an individual’s immune phenotype (Navara and Mendonca, 2008).

Of the hormones thought to influence immune development, testosterone has a reputation as a potent immunomodulator, and has long been described as a general inhibitor of immune activity (Folstad and Karter, 1992; Roberts et al., 2004). Testosterone exposure can suppress functionality of both B lymphocytes (antibody-producing cells) and T lymphocytes (regulators of cell-mediated immunity) *in vitro* (Cunningham and Gilkeson, 2011). In addition, a variety of vertebrate taxa exhibit decreased immune activity when treated with androgens. For instance, in adult mammals, androgen exposure causes apoptosis in cells of the thymus (the site of T cell maturation), as well as in circulating T cells and macrophages (Olsen and Kovacs, 1996). Studies on testosterone exposure during early development have reported similar effects. In birds, *in ovo* testosterone injections decrease post-hatch T cell concentrations as well as suppress the T cell-mediated response to phytohemagglutinin (PHA), a general irritant and T cell mitogen (Andersson et al., 2004; Groothuis and Ros, 2005; Muller et al., 2005; Navara et al., 2005), though results are not consistent across species (for example, Tschirren et al. 2005). In chickens, individuals from eggs treated with high doses of testosterone propionate show depressed antibody levels, as well as reduced weight of the bursa of Fabricius, an exclusively avian organ responsible for B-cell development and acquired immunity (Glick, 1961; Hirota et al., 1976; Norton and Wira, 1977). Thus, in birds, early administration of testosterone can inhibit several types of immune cells and immune cell generation, and these effects can persist after hatching.

### 1.1 Aromatase and Estradiol

Despite evidence for testosterone-induced immunosuppression, our mechanistic understanding of testosterone’s complex role in immunity remains incomplete. Cytochrome P450 aromatase is an enzyme that converts testosterone to 17β estradiol – all estradiol must be derived either from testosterone through this pathway or from androstenedione via estrone as an intermediate (Simpson et al., 2002). In addition, local conversion of testosterone to estradiol by aromatase is a key step in many signaling pathways purported to be testosterone-dependent. For instance, neural exposure to locally aromatized estradiol in early embryogenesis promotes brain masculinization in mice, and aromatase knockout mice show a marked decrease in characteristic male behavior (Matsumoto et al., 2003). Additionally, in song sparrows, display of male territorial aggression during breeding season is dependent on aromatization (Soma et al., 2000). Experiments such as these suggest that many of the effects of testosterone are not regulated via activation of androgen receptors but instead are mediated by aromatization to estradiol; thus aromatization may also play a crucial role in testosterone-dependent regulation of immunity.

The role of aromatization in immunomodulation has been studied extensively within the biomedical field (Niravath, 2013; Subramanian et al., 2008; Tanriverdi et al., 2003) and the prospect of immunosuppression by estradiol is frequently discussed in the context of xenoestrogens and other endocrine disruptors (Chalubinski and Kowalski, 2006; Markman et al., 2008; Razia et al., 2006). However, this pathway is often overlooked in other contexts, despite the long-standing knowledge that maternal testosterone can induce immunosuppression and acknowledgment of aromatization as a potential fate of testosterone (Owen-Ashley et al., 2004). Like testosterone, estradiol inhibits the T cell mediated response to PHA *in vitro* and causes dose-dependent thymic atrophy in mice (Ablin et al., 1974; Okasha et al., 2001; Rijhsinghani et al., 1996; Staples et al., 1999; Tai et al., 2008; Wyle and Kent, 1977). Similarly, estradiol downregulates B cell lymphopoiesis in humans (Hill et al., 2011). The few existing studies in birds are largely restricted to adult or juveniles, and have found that estradiol tends to depress cell-mediated immunity, although reported effects on humoral immunity have been inconsistent (Owen-Ashley et al., 2004). Treatment of immature broiler chicks with estradiol or testosterone resulted in a decline in leukocytes, lymphocytes, and bursal weight, while administration of dihydrotestosterone (DHT), a non-aromatizable androgen, did not (Al-Afaleq and Homeida, 1998). Similarly, estradiol administration to chicken eggs on day 14 of incubation caused a suppressed antibody response to killed *Brucella abortus* 2-4 weeks after hatching, as well as a bursal weight reduction (Kondo et al., 2004). Thus, there are similarities between the immunomodulatory effects of testosterone and estradiol that may implicate aromatization as a mechanism through which developmental exposure to testosterone acts on the immune system. While the importance of aromatization in immune regulation has been acknowledged in mammals (Martin, 2000) and in avian adults and juveniles, to our knowledge no prior studies have explored the role of aromatization during embryonic immune system development in birds.

The purpose of our study was to determine whether conversion of endogenous testosterone to estradiol is involved in regulating development of immune function in birds. We tested this by manipulating aromatase activity during development by exposing chicken embryos to an aromatase inhibitor, Fadrozole, during a critical window for immune tissue development. We tested two contrasting hypotheses (Fig. 1). The first hypothesis (H1) asserts that the reduction in immune function documented in chickens in response to embryonic testosterone administration is due (at least in part) to aromatization of androgens to estrogens. Under this hypothesis, we predicted that aromatase inhibition *in ovo* would result in elevated immune activity relative to controls. The second hypothesis (H2) asserts that aromatization is not involved in regulating immune activity of developing birds. Therefore, under this hypothesis we predicted that exposure to an aromatase inhibitor a) would not change immune activity, or b) may actually decrease immune activity relative to controls if testosterone is immunosuppressive and inhibition of aromatase causes a buildup of testosterone due to lack of negative feedback in the hypothalamus. Note that our conceptual framework is based on studies in chickens which show that testosterone either inhibits or has no effect on immune function, rather that enhancing it.

**Figure 1.**
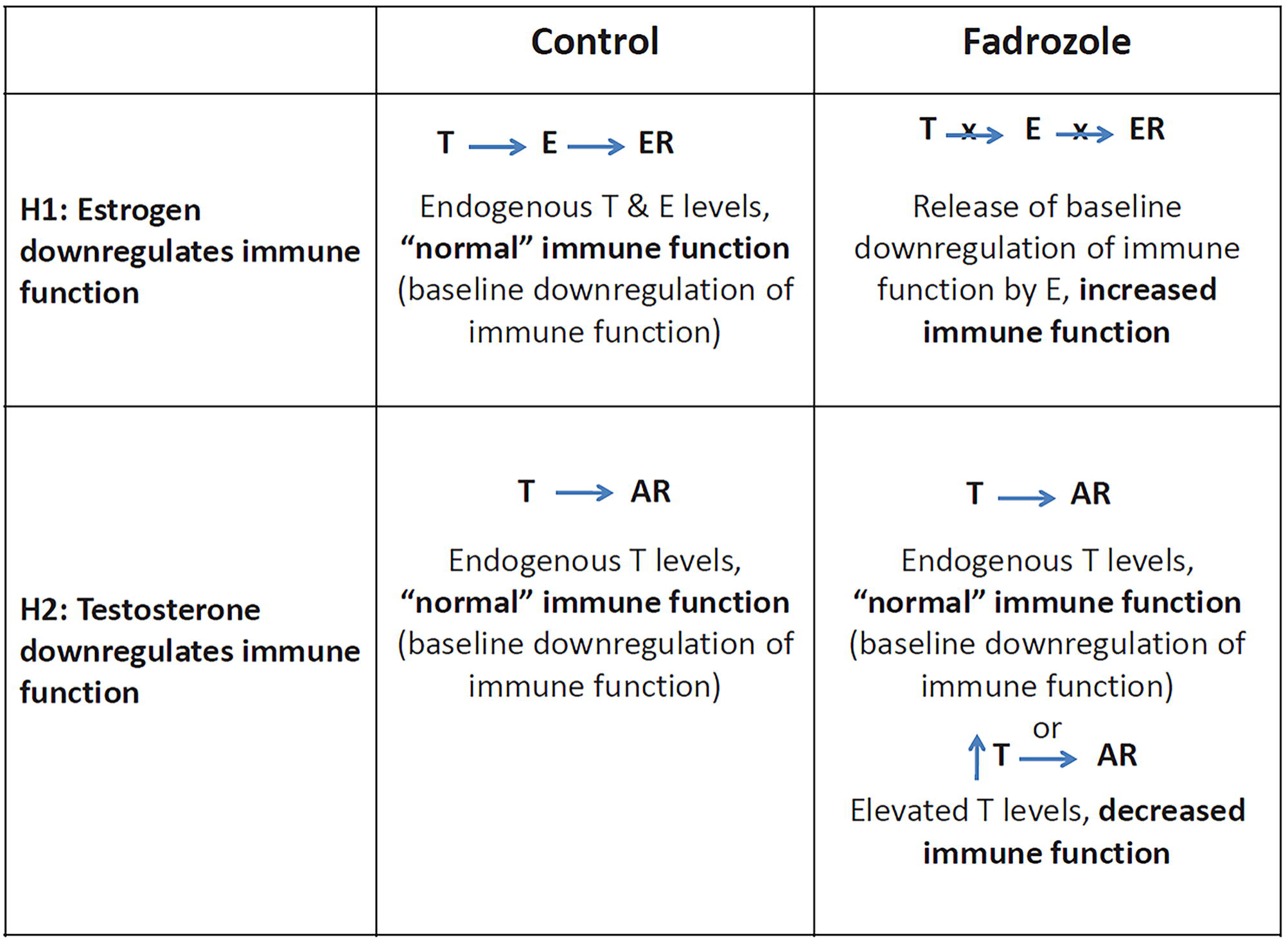
Predictions generated by two contrasting hypotheses (H1 and H2) to explain the immunosuppressive effects of embryonic exposure to testosterone. (T: testosterone, E: estradiol, AR: androgen receptors, ER: estrogen receptors).

## 2. METHODS

### 2.1 Immune traits

We measured several traits representing different branches of the immune system. To measure acquired immunity, plasma IgY antibody levels were assayed at three points during development. IgY is the major immunoglobulin in birds, and is functionally similar to mammalian IgG antibody. Maternal IgY is deposited in egg yolk and is selectively transferred to the embryo via an IgY receptor (FcRY) embedded in the yolk cell membrane (Tesar et al., 2008). This maternally derived IgY acts as the basis of acquired immunity for the developing chick until it is able to produce its own at approximately day 6 post-hatch (Hamal et al., 2006). Because all IgY present before post-hatch day 6 is maternally derived, IgY titers before this point in development reflect uptake or retention of maternal antibodies. After day 6, IgY levels reflect a mixture of maternally derived and endogenous immunoglobulin. By day 14, maternal IgY levels are undetectable and all IgY is synthesized endogenously (Hamal et al., 2006). The mass of the bursa of Fabricius was also recorded as a potential indicator of B lymphocyte generation, as bursal mass and antibody production are correlated (Glick, 1956; Sadler and Glick, 1962).

To assess T cell mediated immunity, swelling in response to PHA injection was measured (Hasselquist and Nilsson, 2012). Additionally, thymic mass was recorded as a coarse indicator of T cell titers, as a correlation between the two measurements has been demonstrated in chickens (Kong et al., 1998).

All procedures were approved by Bucknell University’s IACUC.

### 2.2 Animals and Fadrozole Injection

80 fertile, unincubated White Leghorn domestic chicken (*Gallus gallus*) eggs were obtained from Centurion Poultry in Montadon, PA and incubated at 37.5°C and 50% humidity in an OVA-Easy Advance incubator (Brinsea, Titusville, FL, USA) with a 90-minute turn cycle. On day 13 of incubation, eggs were removed individually from the incubator and 0.1 mg of Fadrozole hydrocholoride (Sigma-Aldrich, St. Louis, MO, USA; F3806) dissolved in 0.1 mL of 0.9% NaCl saline solution or saline only was injected in the air sac using a ½ inch, 27 g needle as previously described (Abinawanto and Saito, 1997; Yang et al., 2008). Egg mass did not differ between treatments (control: 44.9±0.8 g; Fadrozole: 45.3±0.7 g). Fadrozole is a non-steroidal aromatase inhibitor that reduces aromatase activity to near undetectable levels, and thus prevents any endogenous androgens from being converted into estrogens (Yue and Brodie, 1997). The dose of Fadrozole was selected because it eliminates aromatase activity without reducing hatchability in chickens (Yang et al., 2008). To our knowledge, there are no studies that measure the elimination time of a single dose of Fadrozole from an egg. However, administration of 0.1 mg of Fadrozole (the same dose used in this study) to White Leghorn chicken eggs at day 5 of incubation induced full sex reversal in developing females, demonstrating that a single dose during development can have lasting effects on physiology (Abinawanto and Saito, 1997). The timing of Fadrozole injection was selected based on studies showing that lymphoid stem cells have fully colonized the bursal primordium in chickens by day 14 of incubation, and that estrogen receptor expression in embryonic bursa and endogenously-derived plasma testosterone levels in both sexes are at a maximum at this age (Houssaint et al., 1976; Kondo et al., 2004; Woods et al., 1975). These studies suggest that this window of time is critical for estrogen signaling in developing immune tissues, and thus, if estrogen affects immune system development, Fadrozole administration is likely to have pronounced effects during this period.

On day 18 pre-hatch, eggs were transferred to a 1550 Hatcher (GQF, Savannah, GA, USA) at 36.7°C and 60% humidity with no rotation. Chicks hatched on day 21 of incubation. Hatchlings were sexed based on feather structure (chickens of this strain are feather-sexable), banded in the wing web, weighed, and transferred to a brooder maintained at 35°C. 71 (88.75%) chicks hatched over a period of 32 hours. Chicks that had not hatched by 24 hours following the first round of hatching were excluded from the experiment to minimize variability associated with hatching date, leaving a total sample of 58 chicks (16 control female, 16 control male, 12 Fadrozole female, 14 Fadrozole male). Once all experimental chicks had hatched, they were redistributed such that sex and treatment were balanced across three brooders. Chicks were given food and water ad libitum and kept on a 12:12 L:D photoperiod. Chicks were weighed every three days at 9 AM when the lights were turned on. The temperature was reduced by 3°C each week and maintained constant once it reached 30°C. All data were collected and samples analyzed by investigators blind to treatment.

### 2.3 Blood Samples and PHA Challenge

On day 3, day 13, and day 18 post-hatch, blood samples were drawn from the alar vein, using ½ inch, 26 g needles and capillary tubes. Samples were kept on ice and separated by centrifugation within an hour of collection; plasma was removed and stored at -20°C until use in assays.

On day 14 post-hatch, a subset of chicks (36, distributed evenly across sex and treatment) were injected in the wing web of the unbanded wing with 0.1 mg of PHA (Sigma-Aldrich, St. Louis, MO, USA; L8754) dissolved in 0.06 mL of PBS buffer following web thickness measurement with a digital micrometer according to established protocols (Martin et al., 2006). Three measurements were taken and the average was used in calculations. Immediately following initial wing web thickness measurements, a ½ inch, 27 g needle tip on a 100 μL disposable syringe was used to slowly inject the PHA solution into the site of measurement. Nine chicks from each sex and treatment group were injected. 24 hours later (+/- 1 hr), the site of injection was measured again as described above. PHA injection did not influence any of the other metrics that we measured.

### 2.4 Immune Tissue

On day 18 post-hatch, following a final blood sample collection, chicks were euthanized by decapitation. The bursa of Fabricius and thymus were extracted and weighed, and diagonal tarsus length was measured using a digital caliper (VWR, Radnor, PA, USA). Bursal mass and thymic mass were corrected for body size by dividing by overall body mass (as in Kondo et al. 2004).

### 2.5 Sex Determination

Sex was confirmed both by visual inspection of gonads and by genetic sexing (Abinawanto and Saito, 1997). Genetic sex was determined with PCR and gel electrophoresis using AmplitaqGold PCR master mix (Applied Biosystems, Waltham, MA, USA; L00192), primers AvianSex 2550F (GTTACTGATTCGTCTACGAGA) and AvianSex2718R (ATTGAAATGATCCAGTGCTTG) (Fridolfsson and Ellegren, 1999), and 2.5% high-resolution agarose gels, which show one band for males and two for females. One Fadrozole treated chick demonstrated potential sex-reversal: it was initially identified as male based on feathers and gonads, but showed two bands in both of two rounds of genetic sexing. This chick was treated as female for subsequent analyses, though excluding this individual did not alter outcomes.

### 2.6 IgY Content Analysis

Plasma IgY content was determined using a Chicken IgG ELISA Quantification Set (Bethyl, Montgomery, TX, USA; E30-104-25). Dilutions of 1:40000 were consistently in the center of the standard curve and thus samples were analyzed in duplicate at this dilution ratio. Treatment groups were distributed equally across 5 plates with samples from the same chick kept on the same plate. Inter-assay variation based on an internal standard was 15.6%, while intra-assay variation for duplicate samples was typically in the 0-5% range (µ = 2.7±2.5%).

### 2.7 Plasma Testosterone

A salivary testosterone kit (Salimetrics, State College, PA, USA; 1-2312) was used to quantify testosterone in day 3 and 18 plasma. This kit has been used successfully to quantify plasma testosterone in several avian species, and we validated that serial dilutions of chicken plasma were parallel with the standard curve (Marteinson et al., 2011; Washburn et al., 2007). Samples were run across 3 plates, and samples from the same individual at both ages were run on the same plate to reduce inter-assay variation. Inter-assay variation based on high and low controls were 6.7% and 3.4% respectively, and intra-assay variation for duplicate samples was 5.2±5.1%. Testosterone was not detectable in some plasma samples; for these, we set values to 1 pg/mL, the lower detection limit for the assay, but also ran the statistical analyses without the undetectable samples included. We were unable to detect plasma E2 with the available quantity of plasma.

### 2.8 Statistical Analyses

We conducted all of our analyses in R (version 3.0.2). For all of our analyses, we selected the models that best fit the data using model comparison with Akaike’s Information Criterion corrected for small sample size (AICc) using the *dredge* command in the R package *MuMIn* (Symonds and Moussalli, 2011). When more than one model was within 2 AICc of the top model, we report model-averaged effects across the top models using the *model.avg* command, also in *MuMIn* (Burnham and Anderson, 2004; Posada and Buckley, 2004). However, if only one model was best fit (i.e., no other models within 2 AICc), we only report results from that model. For all analyses, we initially ran linear mixed-effects models (LMEs, using the command *lmer* in the package *lme4*) and assessed the effect of inclusion of a random grouping effect of brooder (i.e., the compartment in which hatchlings were housed – hatchlings were housed in 3 different brooders) by comparing the AICc of the full model containing or lacking this random effect. If the AICc was >2 lower when this random effect was included, we retained it in the full model that was then used in model selection for fixed effects. For all models analyzing response metrics that were estimated more than once within individuals, we included individual identity as a random effect. For models where a random effect was not retained (because individuals were only sampled once, and brooder did not improve model fit), we used generalized linear models (GLMs) instead of LMEs. We checked the fit of all top models by visually examining plotted relationships between predicted values and model residuals, and fixed effects and residuals, as well as by testing for normal distribution of residuals.

To determine if the treatment influenced any of the response metrics (production or uptake of IgY, bursal or thymic mass, post-hatch testosterone, or the PHA response) we conducted separate LMEs with the full model containing treatment (control or Fadrozole treated), sex, body mass, and age (3, 13, or 18 days old) as fixed effects where appropriate. We log-transformed IgY and testosterone concentrations to fit a normal distribution. We corrected bursal mass and thymic mass by dividing by body mass, so we did not include body mass in these models. We also included the interactions of treatment by age and treatment by sex. If Fadrozole treatment influenced any of the response variables, the treatment main effect or treatment by age interaction effect would be significant.

We also conducted an additional analysis to determine if treatment with Fadrozole influenced the efficiency of IgY production by the bursa. We used an LME with log-transformed IgY measured at 18 days old as the dependent variable. The full model included treatment, size-corrected bursa (as described above), sex, and body mass, as well as the interactions of bursa by treatment, bursa by sex, treatment by sex, and treatment by sex by bursa as fixed effects. If treatment influenced the production of IgY by the bursa, we would predict that the relationship between bursa mass and IgY would differ for Fadrozole-treated birds (i.e., a significant bursa by treatment interaction effect).

Finally, to determine if treatment with Fadrozole influenced the relationship between testosterone and thymic mass, we conducted an LME with size-corrected thymic mass as the dependent variable. The full model included treatment, log-transformed testosterone (measured at 18 days old), sex, and all possible 2-way interactions and the 3-way interaction as fixed effects. If Fadrozole treatment influences the effect of testosterone on thymic mass, we would predict that the relationship between testosterone and thymic mass would differ for Fadrozole-treated birds (i.e., a significant treatment by testosterone effect).

## 3. RESULTS

### 3.1 Hatching & Growth

All chicks hatched within 24 hours of each other. Hatching success did not differ between treatments (90% (36 out of 40 eggs) for control and 85% (34 out of 40 eggs) for Fadrozole chicks; (χ^2^(1, N = 80) = 0.125, p = 0.723). Fadrozole treatment did not affect hatch mass (F_1,56_ = 0.363, p = 0.549), body mass gain (F_1,56_ = 0.209, p = 0.65), or day 18 tarsus length (F_1,55_ = 0.63, p = 0.431).

### 3.2 Plasma IgY Concentrations

Inclusion of brooder (σ^2^ = 0.002±0.04; all random effects reported as estimated variance ± standard deviation) as a random effect improved model fit, so model comparison was conducted with an LME including both brooder and individual identity (σ^2^ = 0.004±0.06) as random effects (because IgY was estimated in the same individuals three times). The 6 best-fit models for explaining variation in IgY concentrations retained age of the individual, treatment, body mass, sex, and an interaction of treatment by age in various combinations (Table 1). In model-averaged estimates, only age, treatment, and the treatment by age interaction were significantly related to IgY concentrations. Overall, IgY declined with age (β = -0.054, *p* < 0.001) and was lower in Fadrozole-treated birds versus controls (β = -0.071, *p* = 0.018), but this difference diminished with age (β = 0.004, *p* = 0.037; Fig. 2).

**Figure 2.**
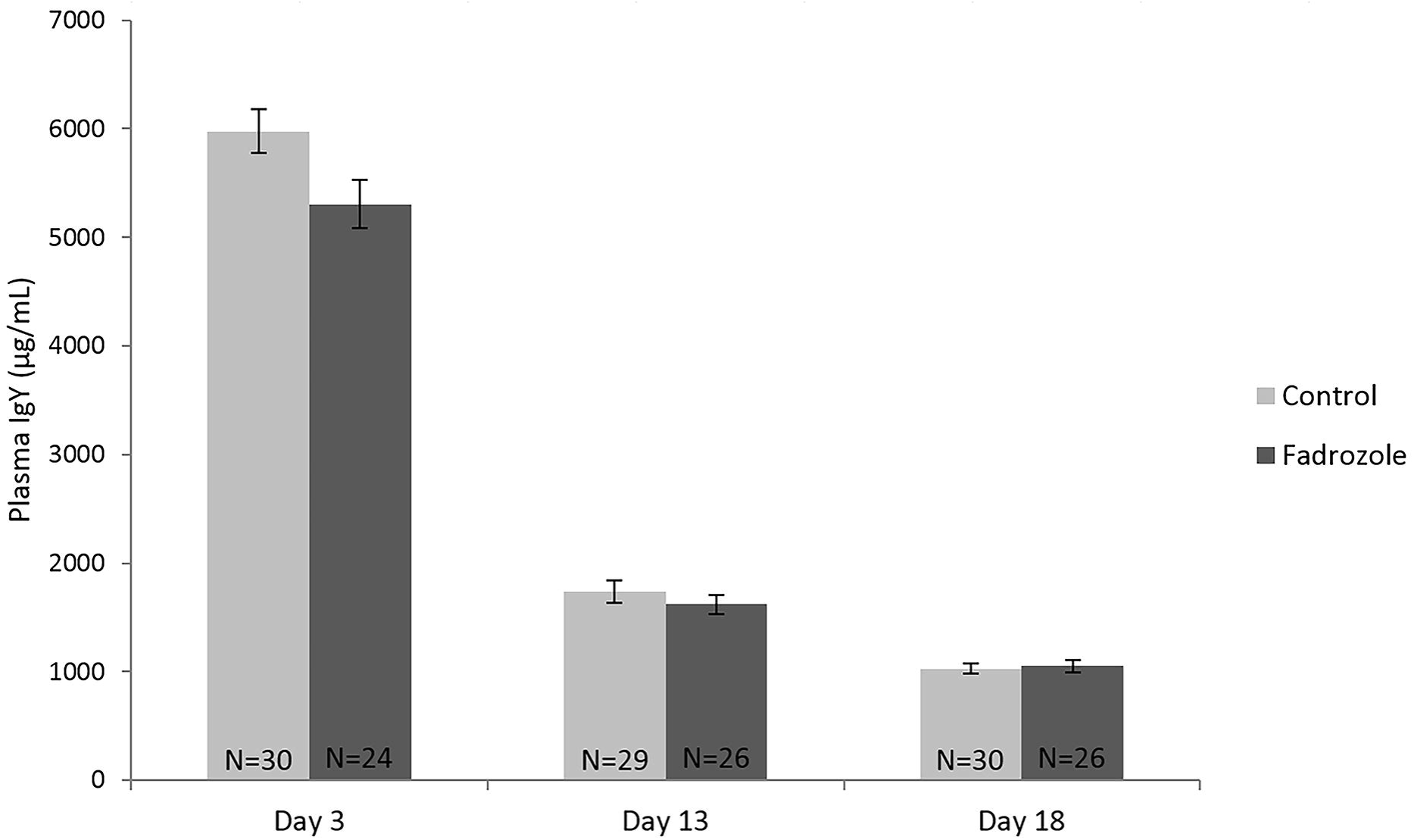
Plasma IgY antibody concentrations for control and Fadrozole chicks (mean±SEM).

**Table 1.** Predictors of IgY concentrations estimated using model averaging for the 6 best-fit models (all within 2 AICc of the top model).

| Fixed effect | Estimate | SE | *z* | *P* |
| --- | --- | --- | --- | --- |
| Age | -0.054 | 0.004 | 12.44 | <0.001 |
| Treatment | -0.071 | 0.030 | 2.37 | 0.018 |
| Body mass | 0.001 | 0.001 | 1.42 | 0.156 |
| Sex | -0.018 | 0.022 | 0.81 | 0.417 |
| Treatment by age | 0.004 | 0.002 | 2.09 | 0.037 |

### 3.3 Bursal Mass

Inclusion of brooder as a random effect did not improve model fit (σ^2^ < 0.0001±0.00), so a GLM without any random effect was used for model selection. The best-fit model was the null model, but 3 other models were similarly well fit to the data as the null (i.e., within 2 AICc). These models individually included fixed effects of phytohemagglutinin (PHA) status (whether or not the bird was injected with PHA), treatment, or sex. However, in model averaging, none of the effects of these factors were significant predictors of bursa mass (all *p* > 0.3). Thus, treatment with Fadrozole did not affect size-corrected bursal mass.

### 3.4 Thymic Mass

Inclusion of brooder as a random effect did improve model fit (σ^2^ < 0.0001±0.00), so model selection was conducted using an LME with this random effect retained. The 3 best-fit models contained sex, treatment, and their interaction (Table 2). Model averaged effects reveal that, overall, males tended to have lower size-corrected thymic mass (β = -0.001, *p* = 0.083), and that Fadrozole-treated birds tended to have higher size-corrected thymic mass (β = 0.001, *p* = 0.056; Fig. 3). The interaction term was not significant.

**Figure 3.**
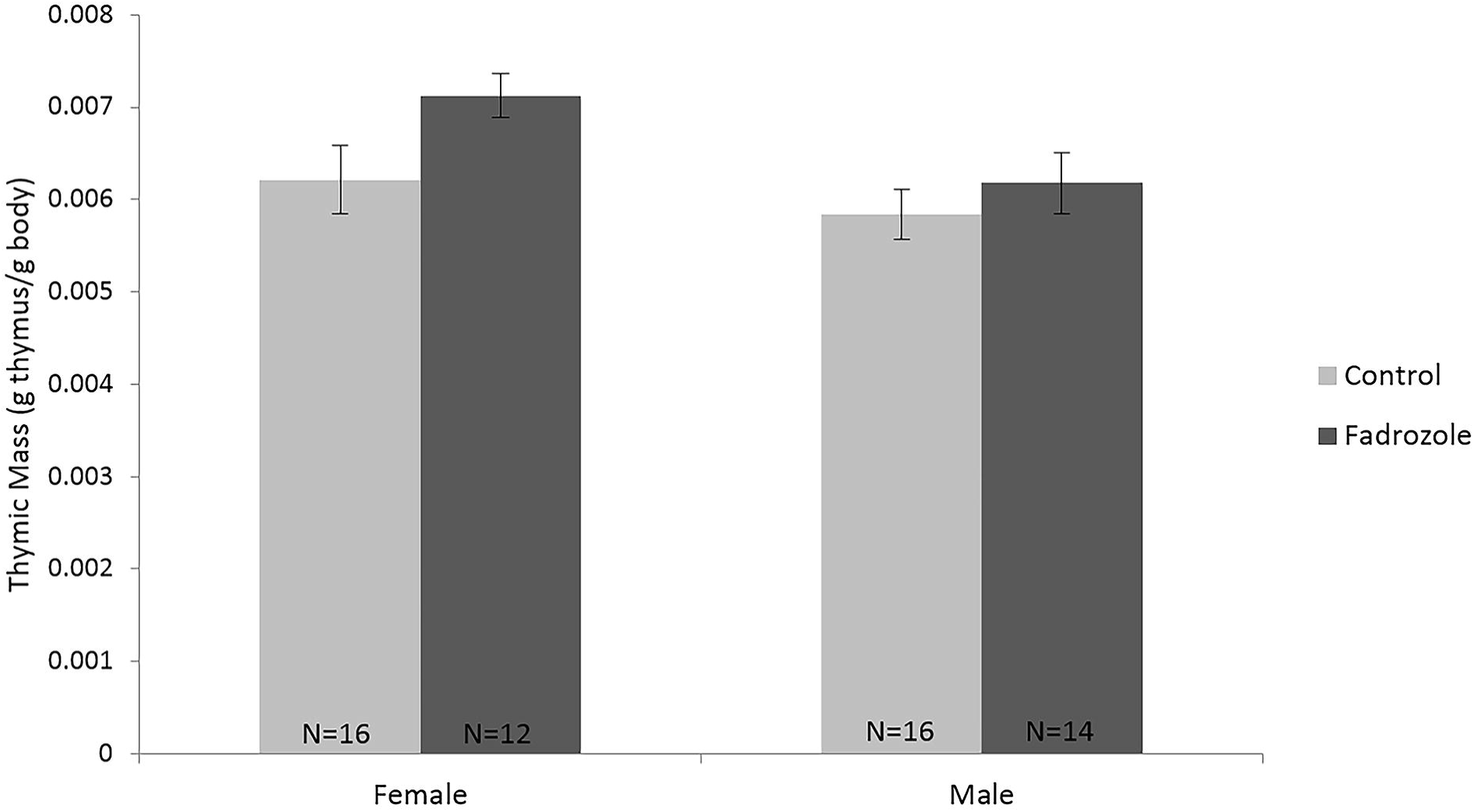
Thymic mass (corrected for body mass) of control and Fadrozole chicks (mean±SEM).

**Table 2.** Predictors of size-corrected thymic mass estimated using model averaging for the 3 best-fit models (all within 2 AICc of the top model).

| Fixed effect | Estimate | SE | $z$ | $P$ |
| --- | --- | --- | --- | --- |
| Treatment | 0.001 | 0.0003 | 1.91 | 0.056 |
| Sex | -0.001 | 0.0003 | 1.73 | 0.083 |
| Sex by treatment | -0.001 | 0.0006 | 0.90 | 0.366 |

### 3.5 Plasma Testosterone Concentration

Inclusion of brooder as a random effect did not improve model fit (σ^2^ = 0.00±0.00), so model selection was conducted using an LME with only individual identity as a random effect (σ^2^ = 0.00±0.00; retained because testosterone was measured twice in each individual). The 3 best fit models retained age, body mass, and sex; treatment was not retained in any of the top models. Model-averaged estimates from the best-fit models indicate that testosterone was higher in males than females (β = 0.856, *p* < 0.001) and declined with age (β = -0. 151, *p* = 0.045). The effect of body mass was not significant (β = 0.007, *p* = 0.202). Results are similar if birds without detectable testosterone concentrations are excluded from the analyses; however, the decline with age is dampened and no longer significant (β = -0.026, *p* = 0.186). Thus, treatment with Fadrozole did not affect circulating testosterone concentrations in 3- or 18-day old birds (Fig. 4).

**Figure 4.**
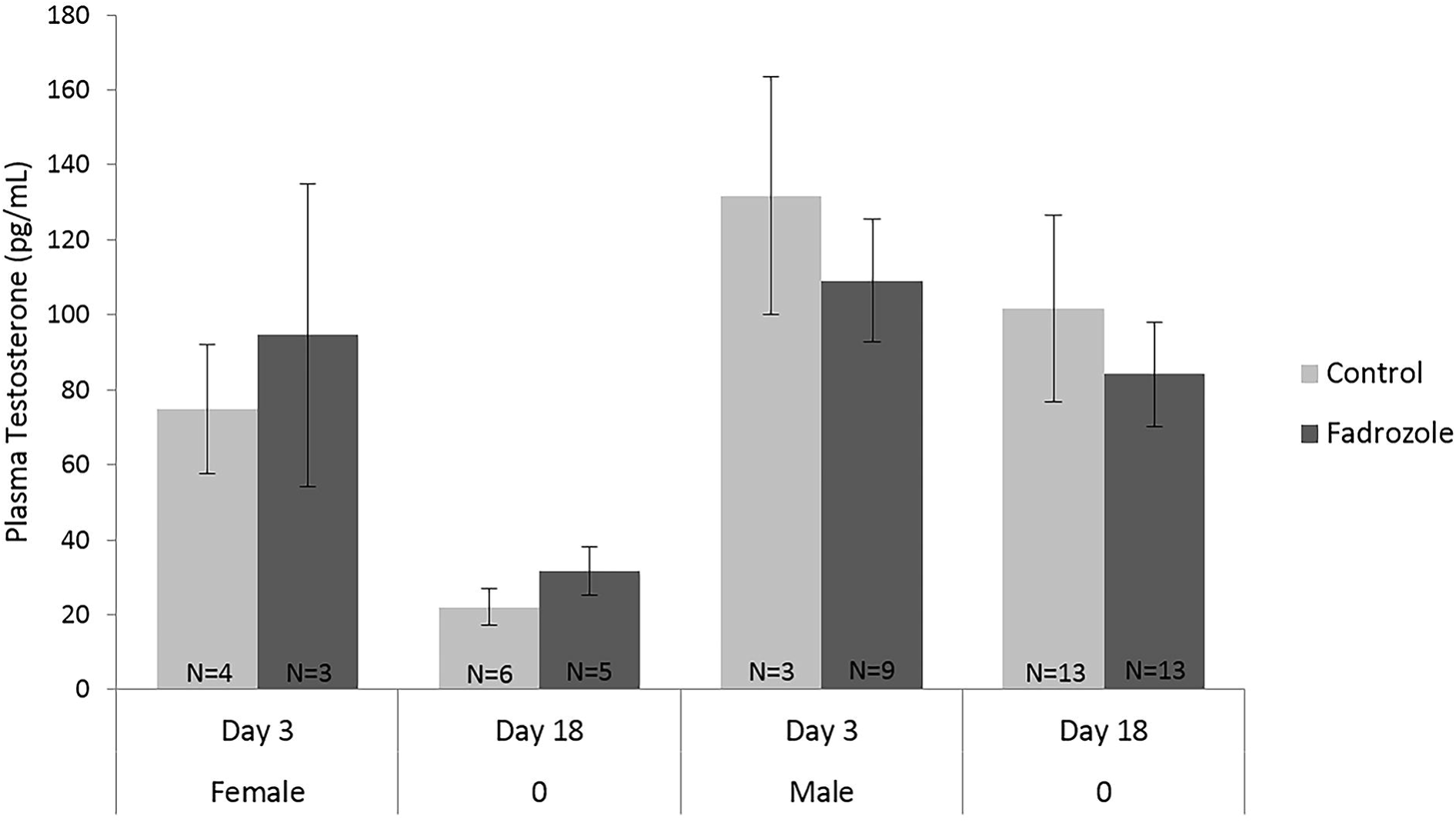
Plasma testosterone levels of control and Fadrozole chicks at day 3 and day 18 post-hatch (mean±SEM). Only individuals with detectable testosterone levels are shown, though relative levels by sex and treatment are unchanged if undetectable samples are included.

### 3.6 IgY-Bursa Relationship

The inclusion of brooder as a random effect (σ^2^ = 0.002±0.04) did not improve model fit, so model selection was conducted with a GLM. The 3 best-fit models retained treatment, size-corrected bursal mass, sex, a sex by bursal mass interaction, a treatment by bursa interaction, and a sex by treatment interaction effect (Table 3). Model averaging revealed both a sex-dependent (β = -104.5, *p* < 0.001) and a treatment-dependent relationship between bursal mass and IgY (β = 67.14, *p* = 0.027). Within females, Fadrozole exposure caused the positive relationship between bursal mass and day 18 IgY to increase in slope, which led to elevated IgY levels for a given bursal mass relative to controls. Conversely, control males exhibited a negative correlation between bursal mass and day 18 IgY, with Fadrozole treatment tending to dampen the relationship (Fig. 5).

**Figure 5.**
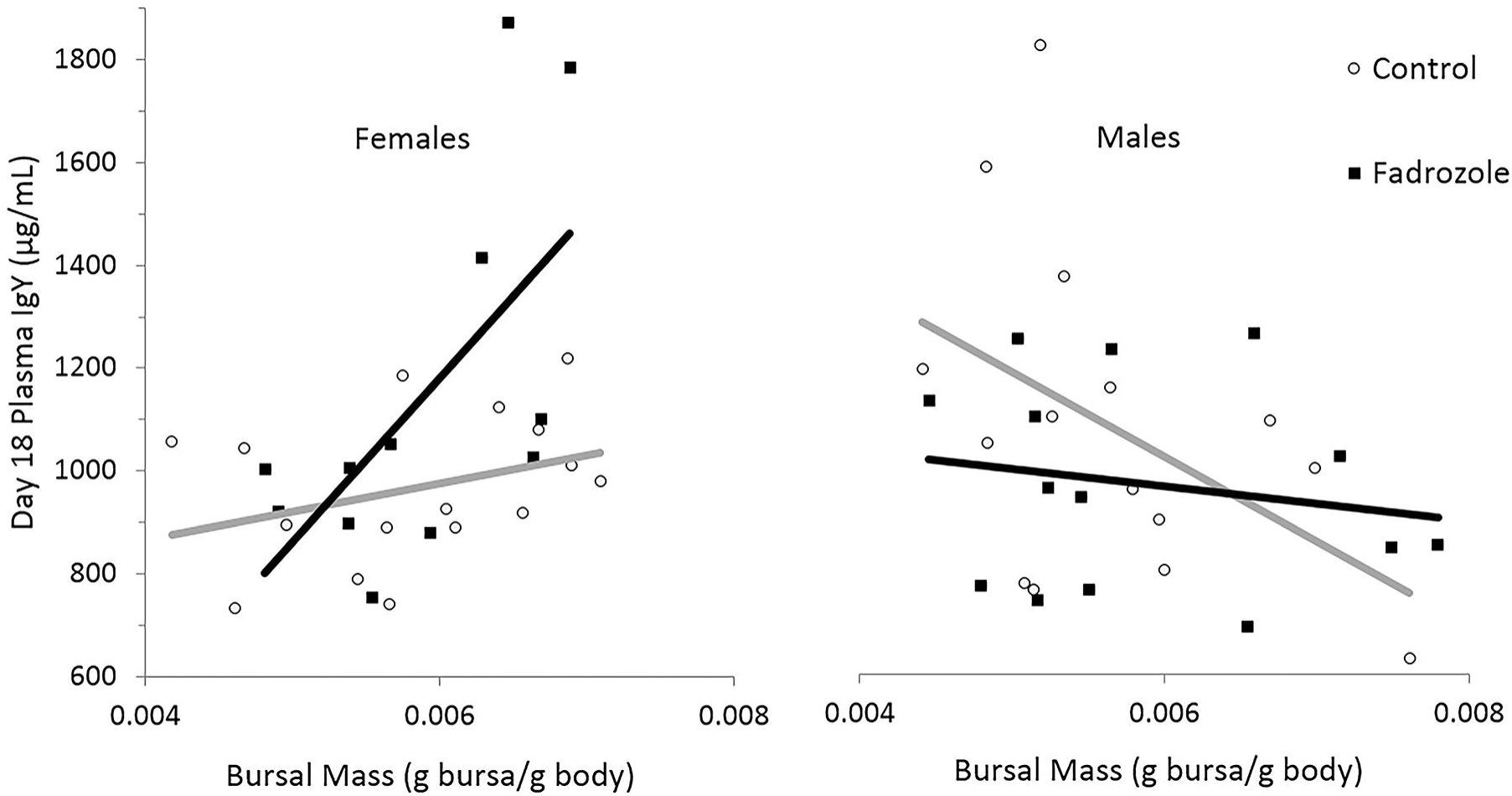
Relationship between bursal mass and day 18 IgY levels by treatment for females (left) and males (right).

**Table 3.** Predictors of day 18 IgY concentrations measured estimated using model averaging for the 3 best-fit models (all within 2 AICc of the top model).

| Fixed effect | Estimate | SE | *z* | *P* |
| --- | --- | --- | --- | --- |
| Treatment | -0.355 | 0.178 | 1.95 | 0.051 |
| Sex | 0.614 | 0.182 | 3.30 | <0.001 |
| Bursa mass | 36.17 | 25.66 | 1.38 | 0.168 |
| Bursa by sex | -104.5 | 30.83 | 3.32 | <0.001 |
| Bursa by treatment | 67.14 | 29.58 | 2.22 | 0.027 |
| Sex by treatment | -0.081 | 0.050 | 1.59 | 0.112 |

### 3.7 Testosterone-thymus relationship

Inclusion of brooder as a random effect did improve model fit (σ^2^ < 0.0001±0.00), so model selection was conducted using a LME with this random effect. The 5 best-fit models retained log-testosterone, sex, treatment, a sex by treatment interaction effect, and a testosterone by treatment interaction effect (Table 4). Model averaging indicated that the interaction of log-testosterone by treatment and sex by treatment were both marginally significant. The majority of females had undetectable testosterone levels at day 18, precluding analysis of the relationship between testosterone and thymic mass. In control males, there was a negative relationship between plasma testosterone and size-corrected thymic mass. In contrast, Fadrozole treated males exhibited a positive relationship between plasma testosterone and size-corrected thymic mass (Fig. 6).

**Figure 6.**
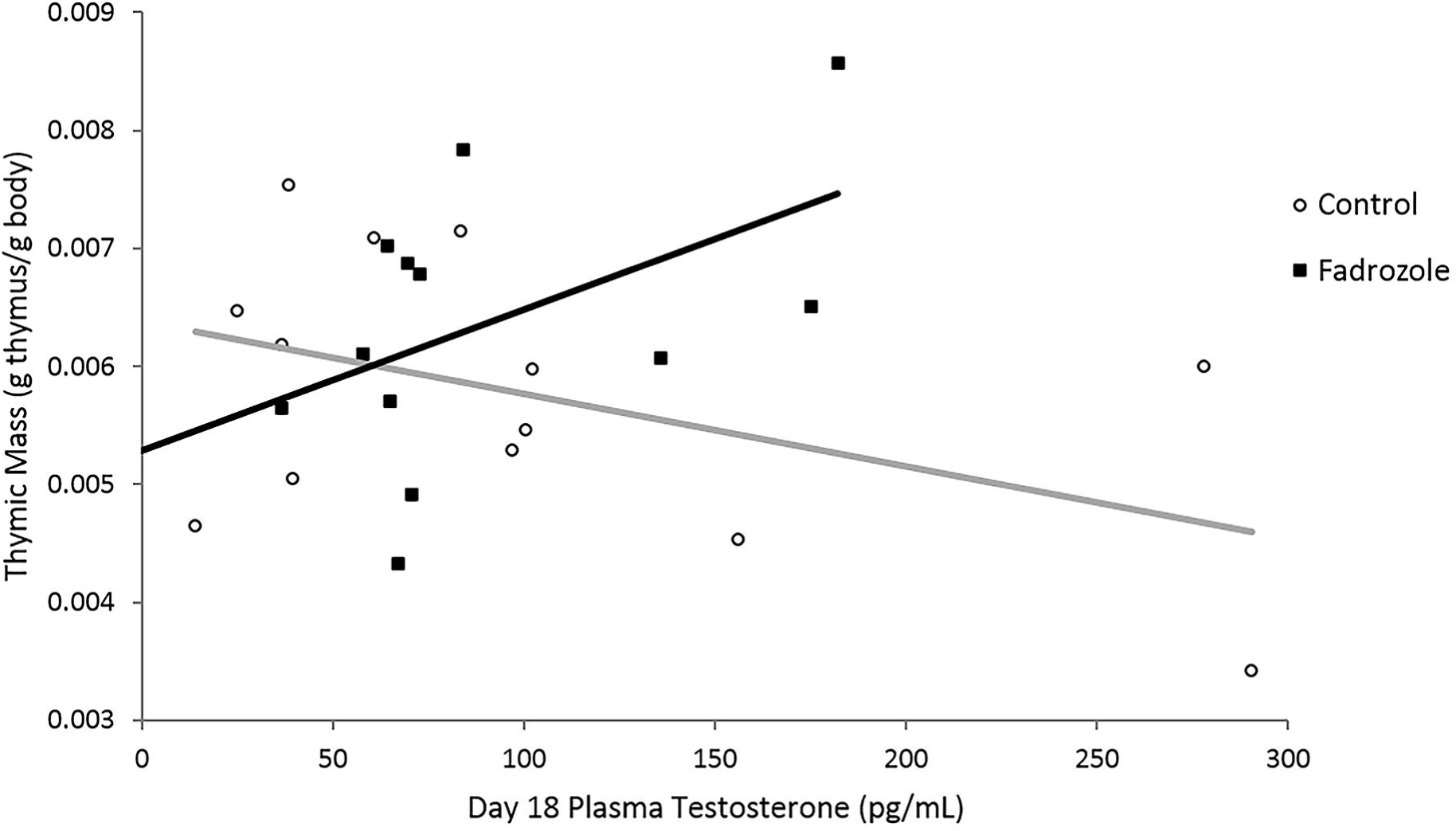
Relationship between day 18 plasma testosterone levels and day 18 thymic mass for males by treatment.

**Table 4.** Predictors of size-corrected thymic mass estimated using model averaging for the 5 best-fit models (all within 2 AICc of the top model).

| Fixed effect | Estimate | SE | *z* | *P* |
| --- | --- | --- | --- | --- |
| Treatment | 0.001 | 0.0004 | 1.40 | 0.161 |
| Sex | -0.001 | 0.0006 | 1.62 | 0.105 |
| Testosterone | 0.0003 | 0.0004 | 0.79 | 0.428 |
| Testosterone by treatment | 0.001 | 0.0006 | 2.03 | 0.043 |
| Sex by treatment | -0.002 | 0.0009 | 1.99 | 0.047 |

### 3.8 Response to PHA

The inclusion of brooder as a random effect (σ^2^ = 0.00±0.00) did not improve model fit, so model selection was conducted with a GLM. The null model was the best-fit model, but 3 other models were similarly well-fit to the data. These 3 models retained sex and body mass in various combinations; treatment was not included in any of the best-fit models. Model averaging across the 4 top models did not reveal any significant effects (all *p* > 0.14). Thus, treatment with Fadrozole did not influence the wing-web swelling response to injection with PHA.

## 4. DISCUSSION

### 4.1 Summary

We show for the first time that prenatal inhibition of aromatase alters aspects of post-natal immunity in birds. Furthermore, because estradiol synthesis was inhibited for a relatively short window of time during embryonic development, the differences observed in chicks suggest that even short-lived changes in embryonic estradiol production can cause persistent changes in aspects of immune function, indicating an organizational role for estrogen on the avian immune system. Aromatase inhibition tended to enhance traits associated with immune function (with the exception of day 3 IgY levels), but did not alter plasma testosterone levels in postnatal animals. Aromatase inhibition increased day 18 thymic mass, and increased day 18 IgY levels for a given bursal mass in females. While thymic mass in control males was negatively correlated with plasma testosterone titers, as expected (Mase and Oishi, 1991), the opposite was true of Fadrozole treated birds, with higher plasma testosterone titers being associated with larger thymuses. Because aromatization was necessary to induce thymic atrophy, our results strongly support hypothesis #1 (Fig. 1), which asserts that conversion of T to estradiol during embryonic development downregulates immune function, and contribute to a growing body of literature suggesting that developmental estrogens may be as important as androgens in shaping the avian immune system (Ahmed, 2000; Al-Afaleq and Homeida, 1998; Owen-Ashley et al., 2004; Roberts et al., 2004).

### 4.2 Endogenous IgY

Fadrozole exposure tended to increase day 18 IgY levels for a given bursal mass in females (Fig. 5). This relationship suggests that estradiol inhibits bursal IgY production and supports the hypothesis that testosterone can modulate immunity via aromatization to estradiol. Because maternal IgY levels become undetectable by day 14 post-hatch, day 18 IgY is entirely endogenous (Hamal et al., 2006). The fact that the effect of Fadrozole on IgY levels varied from day 3 to day 18 also provides evidence that estradiol has different effects on different mechanisms within the immune system. In this case, estradiol may promote uptake of maternal IgY, perhaps via the FcRY receptor expression (see section 4.3) but inhibit endogenous IgY production after hatching (Hamal et al., 2006; Kondo et al., 2004).

### 4.3 Maternal IgY: Day 3 Post-Hatch

Day 3 IgY concentration was the only immune measure that was reduced by Fadrozole exposure (Fig. 2). As IgY is not synthesized endogenously until post-hatch day 6 in chickens, the IgY titers observed at day 3 represent maternal antibodies that have been retained in the developing chick (Hamal et al., 2006). Maternal IgY is transferred from the yolk to the embryo via a selective receptor (FcRY) embedded within the yolk membrane (Tesar et al., 2008). The observed decrease in IgY levels in response to developmental aromatase inhibition can therefore be explained by either a change in uptake (mediated either directly via alteration of FcRY expression or activity, or indirectly through some other means) or by upregulation of IgY catabolism or excretion. We are not able to distinguish between these possibilities in this study, but to our knowledge, this is the first study to demonstrate that inhibition of aromatase alters uptake or retention of maternal IgY antibody. If expression or activity of receptors for maternal antibodies can be directly influenced by hormone exposure during prenatal development, it suggests a potential interaction among different maternal effects (endocrine versus immune). Variation in IgY uptake could have far-reaching effects on the immune system because maternal IgY provides the basis of acquired immunity until well after hatching (Hamal et al., 2006). Thus variation in IgY uptake could introduce variation in the maturing immune system and in susceptibility to early infection.

### 4.4 Thymic Mass

Fadrozole exposure tended to increase body mass-corrected thymic mass of 18 day old chicks (by about 15% in females; Fig. 3), suggesting that endogenous estradiol inhibits thymic growth for a given body size. Our results corroborate previous studies in mammals showing that the inhibitory actions of testosterone on thymic growth are mediated in part by conversion to estradiol (Greenstein et al., 1988), and studies in amphibians showing that estradiol treatment induces apoptosis of thymocytes (Quinde, 2014). It is difficult to decipher the fitness implications of a difference in thymic mass, because the size of the thymus does not always correspond with T cell populations (Hasselbalch et al., 1997). However, a change in thymic mass in response to inhibition of pre-natal estradiol production represents a potentially important alteration in immunological state.

### 4.5 PHA Challenge

Interestingly, Fadrozole exposure had no effect on response to PHA. The PHA response challenge has long been used as proxy for T cell mediated immunity, and has become one of the most widely used ecoimmunological assays due to its simplicity and cost efficiency (Martin et al., 2006). If, then, the PHA response is truly a measure of T cell activity, it raises the question of why an increase in thymic mass in response to aromatase inhibition did not result in a corresponding increase in PHA swelling response. However, the mechanics of PHA-induced inflammation generate a complex relationship between T cell activity and the PHA response. While PHA does indeed induce T lymphocyte proliferation, the downstream inflammatory response involves a host of other cells as well as regulatory factors, and it is ultimately macrophages, heterophils, and basophils that are the direct agents of swelling (McCorkle et al., 1980; Stadecker et al., 1977; Tella et al., 2008). For this reason, it has been suggested that the PHA challenge assay be considered a “multifaceted index of cutaneous immune activity” more so than a direct measure of T cell mediated immunocompetence (Martin et al., 2006). Thus the lack of correlation between thymic mass and PHA response may not be surprising in the wake of the recent reevaluation of the PHA challenge and its implications (Hasselquist and Nilsson, 2012; Kennedy and Nager, 2006; Martin et al., 2006; Tella et al., 2008).

### 4.6 Plasma testosterone

The lack of differences among treatments in day 3 and day 18 plasma testosterone levels suggests that the changes observed were mediated by organizational effects allowing for long-term alteration of physiology despite transient exposure to Fadrozole (testosterone should be elevated in Fadrozole treated chicks if it were still inhibiting aromatase activity at those ages) (Sechman et al., 2003). Because embryonic testosterone and estradiol levels were not quantified, it is not possible to definitively identify whether the effects of aromatase inhibition were mediated by a transient prenatal increase in testosterone concentration (that was no longer detectable by day 3 post-hatch; Fig. 4) or a decrease in prenatal estradiol concentrations. This limitation is difficult to circumvent given the mechanism of negative feedback for testosterone, because negative regulation of testosterone production is mediated in the hypothalamus by locally-aromatized estradiol, rather than testosterone itself. Therefore, by inhibiting estradiol production, we also inherently reduce negative feedback on testosterone manufacture in the hypothalamic–pituitary–gonadal axis (Hayes et al., 2001). However, it seems unlikely that the elevated immune function in chicks from Fadrozole treated eggs was driven by an increase in developmental testosterone exposure, since studies that manipulate *in ovo* testosterone exposure in chickens consistently report inhibition of immune function (Glick, 1956, 1961; Hillgarth and Wingfield, 1997; Hirota et al., 1976; Navara and Mendonca, 2008; Norton and Wira, 1977). In addition, studies in rats provide precedent for a direct immumodulatory role of estradiol.

In one experiment, aging and young adult rats were orchidectomized and surgically implanted with testosterone or testosterone and the aromatase inhibitor 1,4,6-androstatriene-3,17-dione (ATD). The thymus naturally degenerates with age in rats, and the effect was accelerated in the testosterone only treatment. However, the testosterone + aromatase inhibitor treatment caused a restoration of the thymus, and even enlarged the gland in young intact rats, indicating that testosterone-dependent thymic degeneration is contingent on aromatization (Greenstein et al., 1992). Thus, our results, in conjunction with previous studies, strongly implicate aromatization as a pathway through which prenatal exposure to testosterone modulates immunity. If this is the case, then our current understanding of testosterone’s role in avian immunity is incomplete. Manipulation of testosterone either during development or in adulthood is a standard method for studying the role of androgens, yet without accounting for aromatization, it is impossible to distinguish between the effects of testosterone and those of one of its active metabolites, estradiol.

### 4.7 Conclusion

In the present study, we demonstrate that developmental aromatase inhibition results in elevated immune indicators in chickens. Combined with the long-standing observation that exposure to both testosterone, and its downstream metabolite estradiol, generally results in suppression of immune metrics, our results suggest that aromatization plays a crucial role in testosterone-mediated immunosuppression.

Experimental ablation alone is not sufficient to fully elucidate a hormone’s role in a physiological process (Zera et al., 2007), and a more integrative investigation, including ablation and replacement experiments, molecular studies of estrogen receptors, and fitness studies, will be needed to disentangle the relative contribution of estradiol to immune function. However, the present study provides valuable groundwork towards this end. Considering the paucity of data regarding estradiol-mediated immunosuppression in the context of physiological and behavioral ecology relative to its precursor, testosterone, future investigations into endocrine-immune interactions may benefit from increased investigation into the potential role of aromatization.

## 5. ACKNOWLEDGEMENTS

All procedures were approved by Bucknell University’s IACUC. We are grateful to K. McAvoy, F. Sanjana, K. Fisher, and T. Kenny for assisting in experimental procedures and animal maintenance, K. Field and M. Moore for assistance with immune techniques, C. Rhone for overseeing animal maintenance and Centurion Poultry in Lewisburg, PA for their donation of eggs. Several anonymous reviewers provided valuable feedback on the manuscript. This research was funded by the Bucknell University Biology department and a Robert P. Vidinghoff Memorial Summer Internship from Bucknell University to J.S.

